# The metabolic cost of overcoming air resistive forces in distance running

**DOI:** 10.1101/2021.04.27.441608

**Authors:** Edson Soares da Silva, Rodger Kram, Wouter Hoogkamer

## Abstract

We lack a mechanistic understanding of the relationship between aerodynamic drag forces and metabolic power during running. Further, the energetic and time savings possible from reducing aerodynamic drag (drafting) are still unclear due to the different methods previously assumed for converting from force reductions to metabolic power savings. Here, we quantified how small horizontal impeding forces (equivalent to aerodynamic forces) affect metabolic power and ground reaction forces over a range of velocities in competitive runners. In three sessions, 12 runners completed six 5-minute trials with 5 minutes of recovery in-between. We tested one velocity per session (12, 14 and 16 km/h), at three horizontal impeding force conditions (0, 4 and 8 N). On average, metabolic power increased by 6.13% per 1% body weight of horizontal impeding force but varied considerably between individuals. With greater horizontal impeding force, braking impulses decreased while propulsive impulses increased (p < 0.001). Across running velocities, the changes in braking and propulsive impulses with greater impeding force were correlated (r = -0.97; p < 0.001), but were not related to individual changes in metabolic power. We estimate that at ∼2-hour marathon pace, overcoming air resistive force comprises 8.52% of the gross metabolic power on average.

## INTRODUCTION

In 2018, Eliud Kipchoge ran the official world marathon record of 2:01:39 in Berlin and then in 2019, he ran a 1:59:40 marathon in Vienna. One major difference between the two marathons was air resistance. In Berlin, Kipchoge ran the last 17 km without any aerodynamic drafting. In contrast, in Vienna, for the first 41 km, Kipchoge had interchanging teams of runners specifically positioned to provide substantial drafting. Drafting is the practice of having runners positioned in front of a designated runner so as to reduce the air resistance and hence the metabolic power requirement of the designated runner (1). Drafting allows the designated athlete to run at a faster speed with the same sustainable metabolic power (2), and thereby enhances performance (3). Several recent studies have performed calculations and computer simulations regarding the aerodynamics and energetics of drafting (4, 5, 6, 7). Some relied on Pugh’s measurements of oxygen uptake in a single subject running in a wind tunnel on a treadmill, drafting behind another runner (8). Others used different methods for converting from aerodynamic force to the metabolic cost of running and hence running performance. Because of uncertainty in these previous studies, our goal was to develop an empirical equation for the metabolic power required to overcome aerodynamic drag forces and hence infer the performance effects. Further, we sought to understand the mechanism behind the metabolic savings.

Classic and modern methods yield remarkably similar estimates for the aerodynamic drag force acting on an elite runner and the corresponding mechanical power. For a runner of Kipchoge’s size running solo at 5.86 m/s (2-hour marathon pace), in still air, the equations of Hill (9) predict a force of 8.06 N (see Electronic Supplementary Material Appendix S1) whereas Beves & Ferguson (4) arrived at a value of 6.6 N using modern computational fluid dynamic (CFD) modelling. Polidori *et al*. (7) also used CFD and presented a value of 7.77 N for a similarly sized athlete, Kenenisa Bekele, running solo at 5.75 m/s. A force of 7 N is just over 1% of these runners’ body weights (see Electronic Supplementary Material Appendix S1 for details). The product of force and running velocity equals the mechanical power required to overcome aerodynamic drag. Thus, the corresponding values for mechanical power at 5.86 m/s are also very similar: 47.2 W for Hill (9), 38.6 W for Beves & Ferguson (4) and 44.7 W for Polidori *et al*. (7). However, because those three studies each used a different and dubious method of converting to metabolic power, they surmised that overcoming aerodynamic drag comprises 3% (9), 9-10% (4) and 2.8% (7) of the metabolic cost of running (see Electronic Supplementary Material Appendix S1 for details).

A second and much more direct method of estimating the metabolic cost of overcoming air resistance involves having a runner on a treadmill in a wind tunnel with the fans turned off and then on with the wind tunnel air velocity matching the treadmill belt speed. Pugh (8) pioneered this approach and estimated that at 6 m/s, 7.5% of the gross oxygen uptake rate is devoted to overcoming aerodynamic drag. Later, using the same method (and wind tunnel), Davies (10) estimated that air resistance accounted for 2% of the gross metabolic rate at 5 m/s and 4% at 6 m/s. While the wind tunnel studies have provided valuable insights, they were performed with very small sample sizes (n = 1, (8) and n = 3 (10) and their estimates vary by 2-fold. Thus prior to the present study, we lacked an understanding of the inter-individual variation in responses.

A third conceptually very similar approach compares the metabolic power required during treadmill (i.e. no air resistance) vs. overground running. This method reveals no/little effect of air resistance at slower running velocities (11, 12, 13). However, at 5 m/s, Jones & Doust (13) found overground running was 7% more expensive. Similarly, Pugh (2) found that at 6 m/s, overground running required 9.2% greater oxygen uptake (n = 7).

Finally, scientists have directly measured the increase in metabolic power consumption when horizontal impeding forces are applied to the waists of runners on a treadmill. We interpolated the results of each of these studies to quantify the per cent increase in metabolic power in response to an impeding force of 1% of body weight (BW), see Electronic Supplementary Material Appendix S2 for details). At a running velocity of 3.6 m/s, Lloyd & Zacks (14) found an average 7.9% increase in metabolic power per 1% BW impeding force (n = 3). Soon thereafter, Zacks (15) found a similar average of 7.9% increase per 1% BW impeding force at running velocities between 3.88 and 7.72 m/s but with individual responses ranging from 5.3 to 10.6% (n = 3). However, for running at 3.3 m/s, the results of Chang & Kram (16) reveal an average increase in metabolic power of only 4.7% in response to a 1% BW impeding force (n = 10).

Given the variety of different experimental approaches, the variable findings, the small sample sizes and the considerable inter-subject variability in metabolic power responses to resistive forces of previous studies, we aimed to more systematically quantify how small impeding forces (comparable to air resistance) affect metabolic power in a larger sample of competitive runners over a range of velocities. We hypothesized that metabolic power would increase linearly with increasing horizontal impeding forces. These data should facilitate more accurate calculations of the effect of altered aerodynamic forces on distance running performance.

## METHODS

### Participants

Twelve male runners (age: 26.1 ± 3.5 years, mass: 66.5 ± 5.6 kg, height 1.79 ± 0.09 m) participated. They all had recently run a sub-32-minute 10 km race or an equivalent performance in another distance-running event. The study was performed in accordance with the ethical standards of the Declaration of Helsinki. Ethics approval was obtained from the University of Colorado Institutional Review Board (Protocol#18 - 0110).

### Experimental Protocol

The study consisted of three data collection sessions. During session 1, the subjects completed a health screening form and then provided informed consent. During all three sessions, we measured height and body mass and thereafter, the subjects warmed-up by running on a custom-built force-instrumented treadmill (17) for 3 minutes at 3.33 m/s (12 km/h) followed by 3 minutes at 3.89 m/s (14 km/h). The subjects then ran six 5-minute trials with 5 minutes recovery in-between. We tested one velocity per session (3.33, 3.89 or 4.44 m/s (16 km/h)), at three horizontal impeding force conditions (0, 4 and 8 N). Subjects ran with each horizontal impeding force condition twice per visit, in a mirrored order, which was counterbalanced and randomly assigned. We averaged the two values for each condition.

### Horizontal Pulling Apparatus

To simulate running with air resistance, we applied small horizontal impeding forces at the waist of the runners, near their center of mass (Figure 1). These forces resulted from a hanging mass that was connected via rubber tubing around pulleys to a waist belt. We used long pieces of low stiffness natural latex rubber to minimize both bouncing of the hanging mass and force fluctuations due to length changes in the rubber tubing resulting from slight anterior-posterior movements of the runner on the treadmill. The rubber tubing first passed under a low-friction pulley that could be positioned vertically to match the height of the subject’s waist, ensuring that the impeding force was horizontal. The tubing then attached to an S-beam force transducer (LCCB-50, OMEGA Engineering, Inc., Norwalk, CT, USA) which measured the pulling force and fluctuations throughout the running stride. Another piece of rubber tubing attached to the force transducer and passed over a second low-friction pulley, positioned approximately 6 m high. Hanging masses of 408 and 815 g applied impeding force of 4 and 8 N, respectively. To counterbalance the weight of the force transducer, we added 305 g of lead to the hanging mass. The rubber tubing dimensions differed for the two resistive force conditions: for 4 N we used 3.2/1.2 mm (outer diameter/inner diameter); for 8 N we used 5.6/1.2 mm. The rubber tubing unstretched lengths also differed such that during the running trials the hanging mass hovered about 0.3 m above the floor.

**Figure 1.**
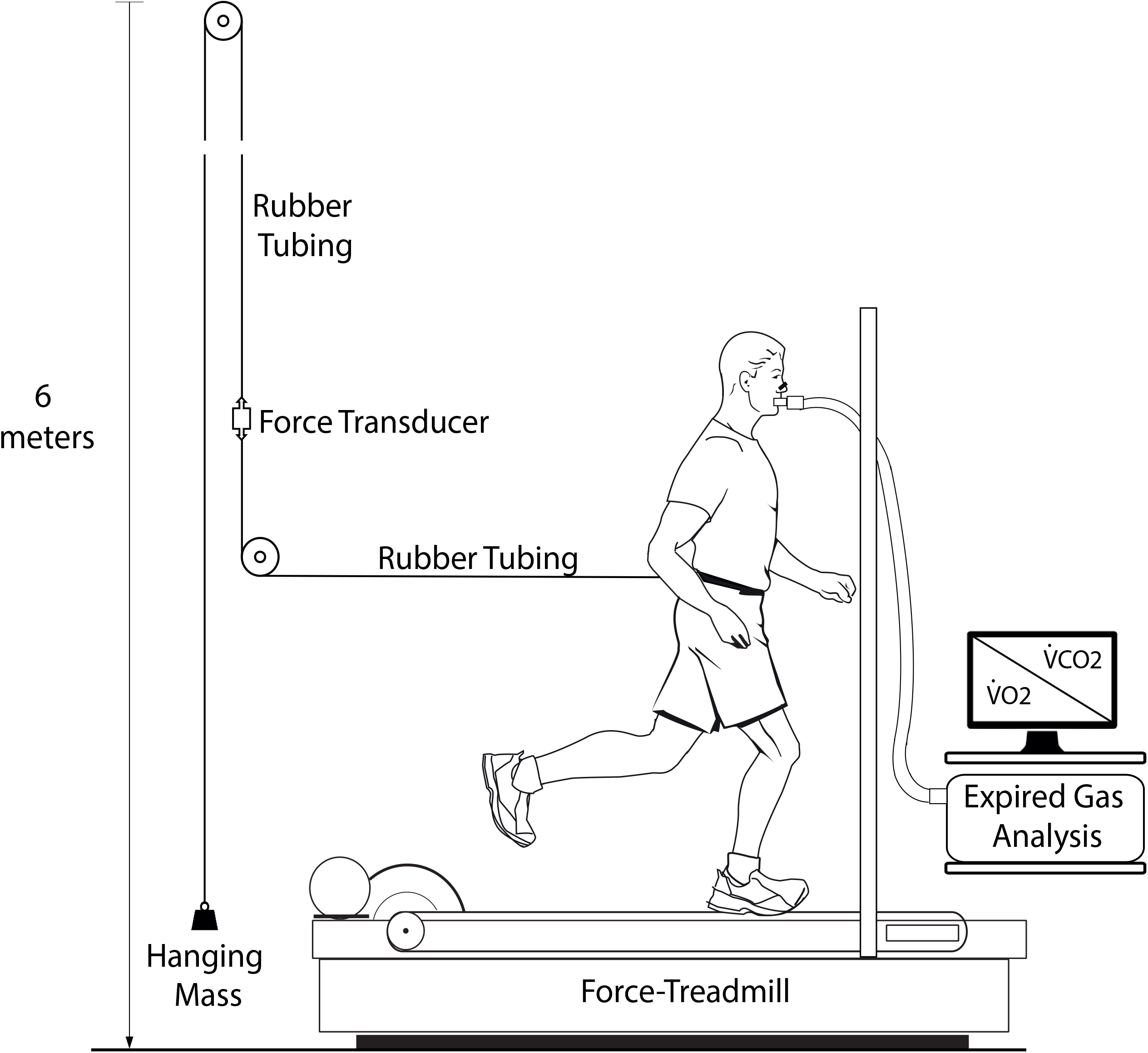
Experimental set-up.

### Metabolic Power Protocol

During each trial, we measured oxygen uptake 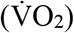 and carbon dioxide production 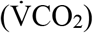 using a breath-by-breath expired air analysis system (True One 2400, Parvo Medics, Salt Lake City, UT, USA) and calculated metabolic power for the last 2 minutes of each trial using the Péronnet & Massicotte (18) equation. Respiratory exchange ratios 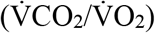 remained < 0.95 for all trials. Body mass was carefully monitored between trials and subjects sipped water to maintain a constant starting body mass for all trials.

### Force Measurements and Analyses

We recorded vertical (F_z_) and anteroposterior (F_y_) ground reaction forces and impeding force fluctuations at a 1000 Hz sampling frequency for 30 seconds during the 2^nd^ and 5^th^ minute using LabView software (National Instruments, Austin, TX, USA). In MATLAB (The MathWorks, Inc., Natick, MA, USA). We filtered the signals (low-pass 4_th_ order Butterworth with cutoff frequency of 25 Hz) and used a 30 N F_z_ threshold to determine touchdown and takeoff events (19). We calculated peak braking and propulsive forces, braking and propulsive impulses (the time integral of force), step frequency and contact time.

### Apparent Mechanical Efficiency

We calculated external mechanical power (W) by multiplying the impeding force (N) by the running velocity (m/s). For each runner, we calculated the “apparent mechanical efficiency” for each impeding force at all three running velocities as the change in external mechanical power (W/kg) from unloaded running divided by the change in metabolic power (W/kg) from unloaded running (14).

### Statistics

We compared metabolic power, temporal and kinetic variables between the three running velocities and the three horizontal impeding force conditions using a two-way ANOVA with repeated measures. When significant main or interaction effects were detected, we performed Bonferroni corrected paired t-tests to determine *post-hoc* which velocity and/or impeding force comparisons differed significantly. We also explored whether inter-individual differences in the increases in metabolic power with impeding force were related to changes in braking or propulsive impulses with impeding force using linear regression analysis. We used traditional levels of significance (α = 0.05 and α_post-hoc_ = 0.0167) and performed analyses with MATLAB.

## RESULTS

Metabolic power was significantly greater at faster running velocities (p < 0.001), and with greater horizontal impeding forces (p < 0.001), with a significant interaction effect (p < 0.001; Figure 2, see Electronic Supplementary Material Appendix S3 for a Table with individual data). The interaction effects were that in response to a specific horizontal impeding force, metabolic power increased more at faster running velocities [12 km/h, 14 km/h and 16 km/h, (p < 0.001)] and at a specific velocity, metabolic power increased more with greater horizontal impeding force [baseline, 4 N and 8 N (p < 0.001)].

**Figure 2.**
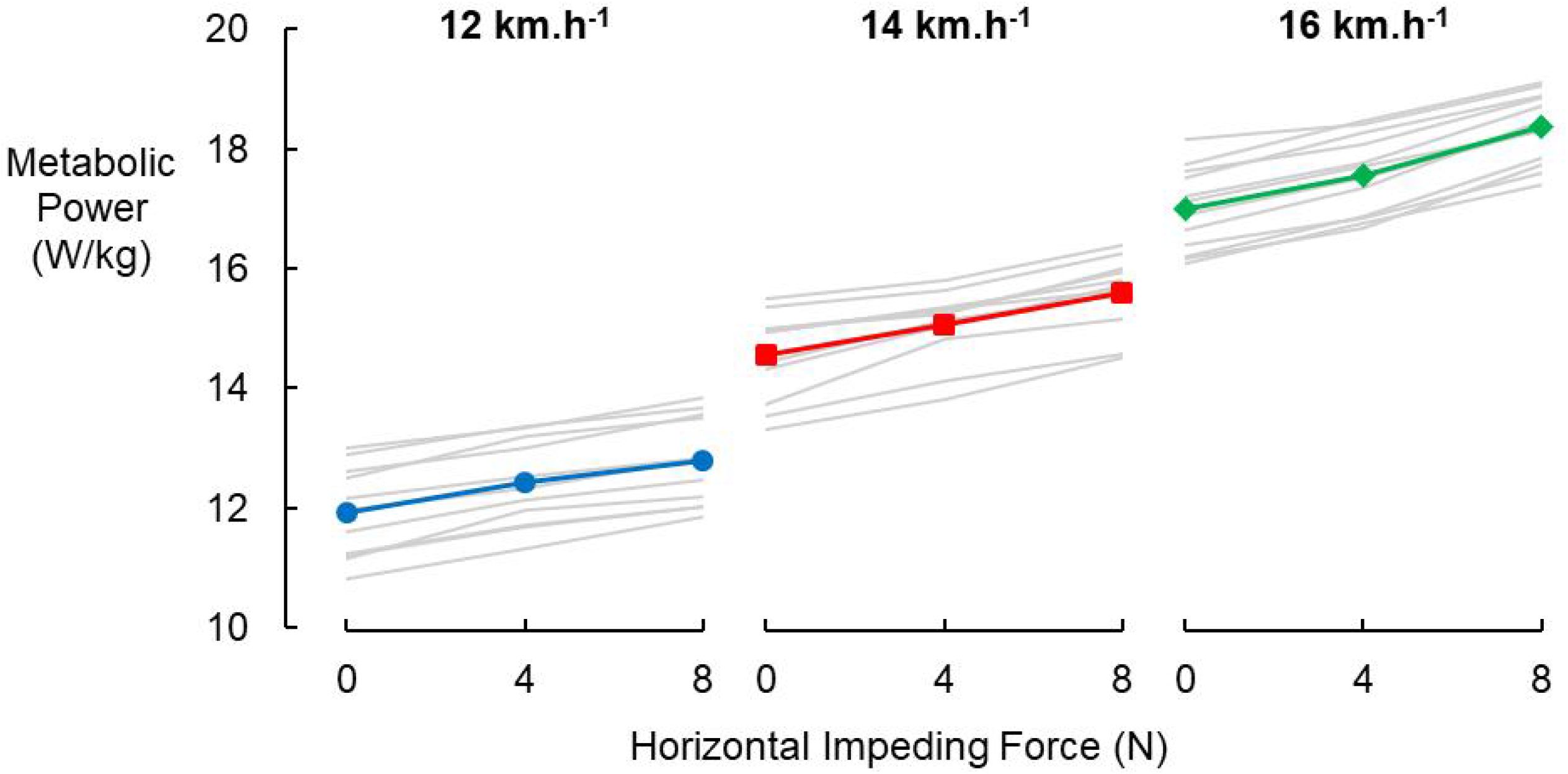
Metabolic power (W/kg) vs. horizontal impeding force (N) for each runner (grey) and the group means (colored symbols) for each of the three velocities tested.

At each velocity, metabolic power increased linearly with increasing horizontal impeding force expressed relative to BW (Figure 3); second order polynomial fitting did not substantially improve r^2^ values. Across runners, the average increase in metabolic rate was 6.13% per 1% BW horizontal impeding force. This was consistent across the three tested running velocities with 6.14, 5.87 and 6.37% slopes for 12, 14 and 16 km/h, respectively. Notably, relative changes in metabolic power with horizontal impeding force varied substantially between individual runners (Figure 4), ranging from 4.75% to 8.14%.

**Figure 3.**
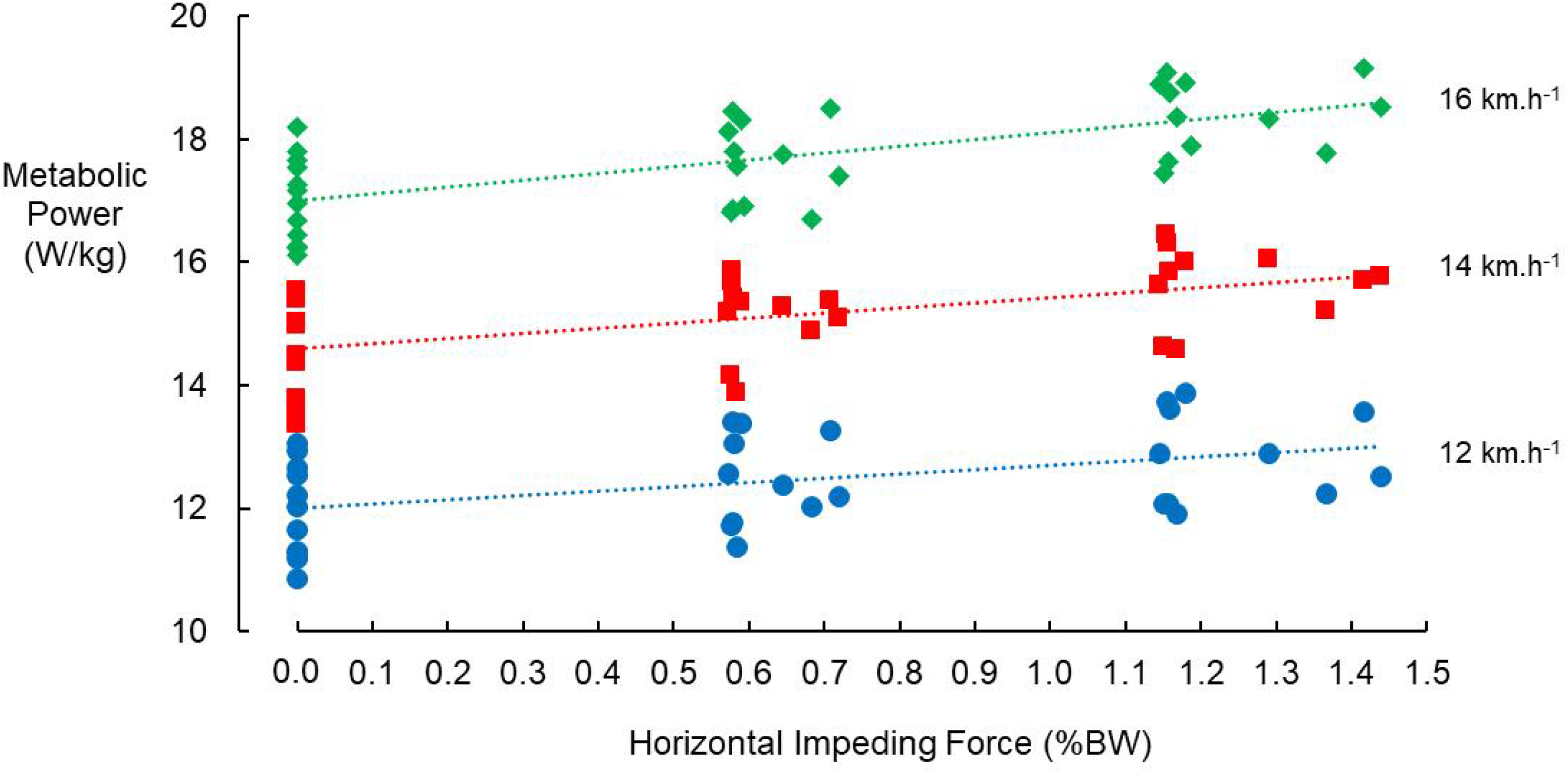
Metabolic power (W/kg) vs. horizontal impeding force (HIF) (% body weight). Blue circles represent individual subjects at 12 km/h, red squares 14 km/h and green diamonds 16 km/h. Dotted lines are the linear best fit regressions at each velocity: at 12 km/h [W/kg = 0.6977 HIF + 11.996 (r^2^ = 0.2004)], at 14 km/h [W/kg = 0.8386 HIF + 14.594 (r^2^ = 0.3193)] and at 16 km/h [W/kg = 1.1048 HIF + 16.993 (r^2^ = 0.4478)].

**Figure 4.**
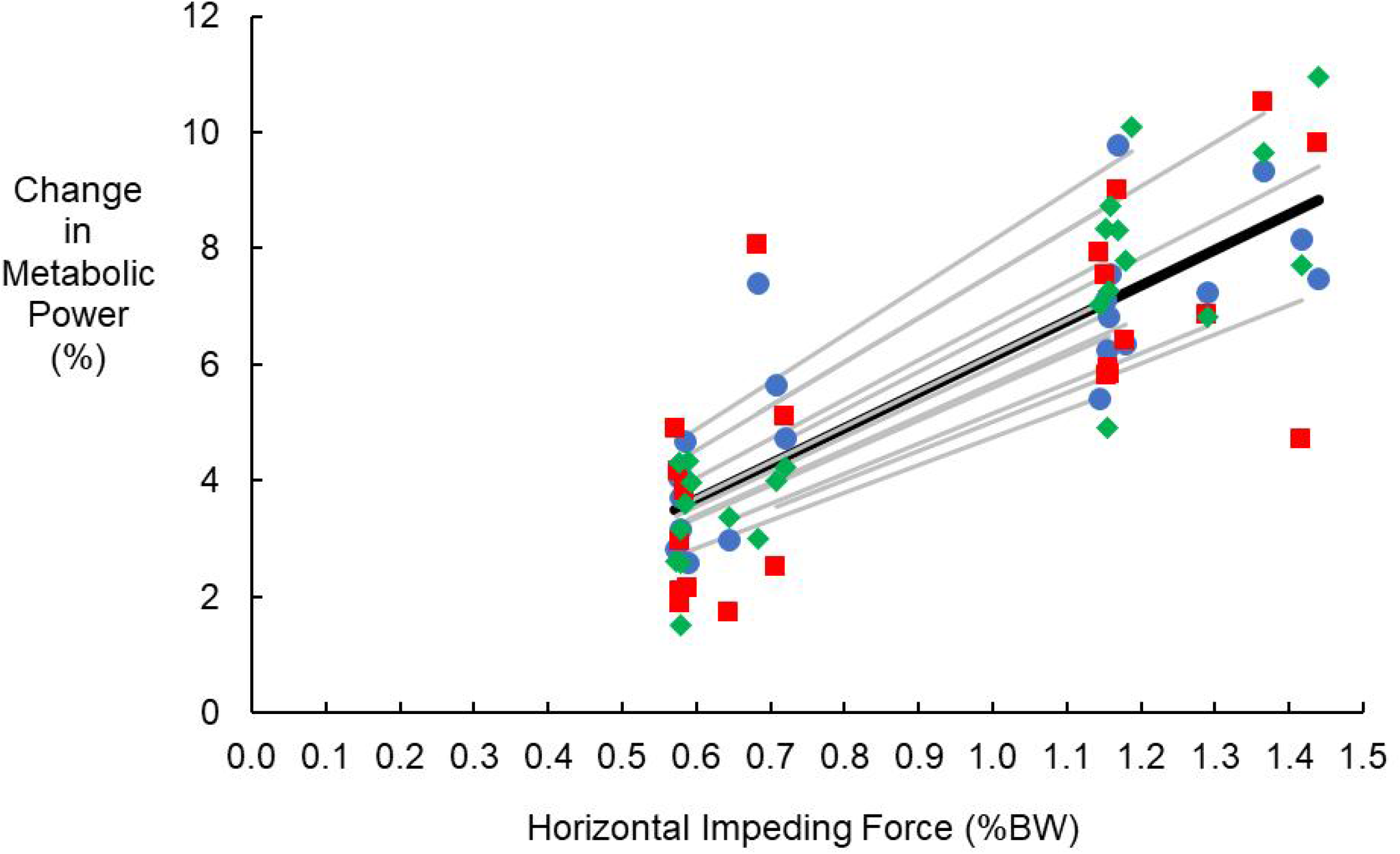
Per cent increase in metabolic power (%) with horizontal impeding force (HIF) (% body weight). Blue circles represent individual subjects at 12 km/h, red squares 14 km/h and green diamonds 16 km/h. For each individual, the best linear fit through the origin is shown in grey. The black line represents the best fit through all the data [% change = 6.13 HIF (r^2^ = 0.68)]. Regressions were forced to go through the origin but zero HIF data points were not included in the regression analysis.

**Figure 5.**
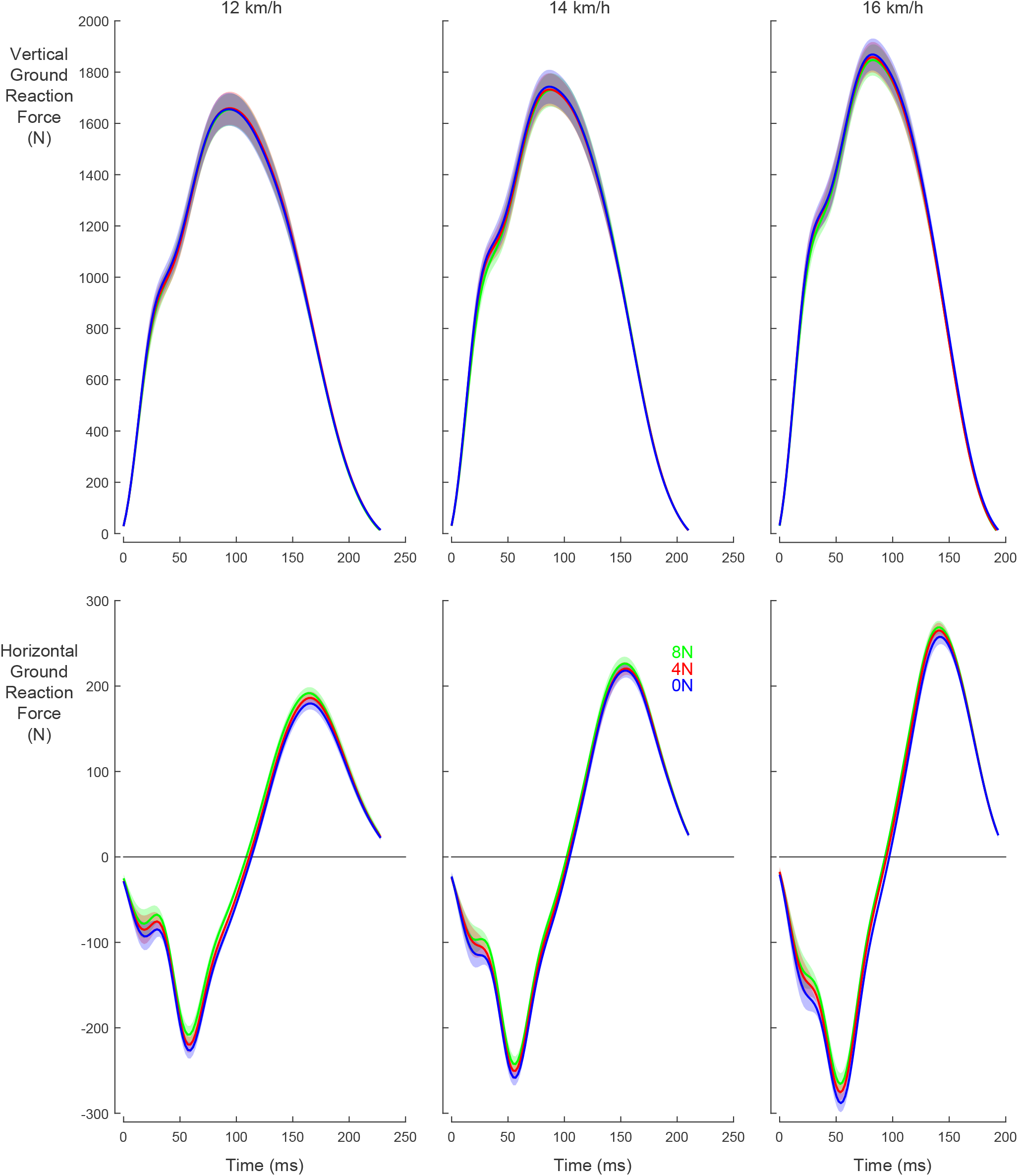
Mean and standard deviation of vertical and anteroposterior ground reaction forces at three running velocities (12, 14 and 16 km/h) and with different horizontal impeding forces (0, 4 and 8N).

Apparent mechanical efficiency was also consistent across the tested velocities (p = 0.401). At the slowest running velocity (12 km/h), apparent mechanical efficiency was 43.6 ± 10.1% from 0 – 4 N and 46.5 ± 5.9% for 0 – 8 N. At the intermediate and fast running velocities, apparent mechanical efficiencies were numerically lower for the stronger impeding forces (14 km/h: 55.2 ± 22.6% from 0 – 4 N and 46.6 ± 11.3% for 0 – 8 N; 16 km/h: 50.1 ± 15.5% from 0 – 4 N and 40.0 ± 7.0% for 0 – 8 N), but these effects were not deemed statistically significant (main effect of impeding force: p = 0.062; interaction effect of velocity by impeding force: p = 0.066).

Step frequency, contact time and duty factor were all independent of horizontal impeding forces (p = 0.061, p = 0.091 and p = 0.786, respectively), but step frequency increased, and contact time and duty factor decreased with running velocity (p = 0.01 0, p < 0.001, and p < 0.001, respectively; Table 2). With increasing horizontal impeding force, braking impulses decreased while propulsive impulses increased (both p < 0.001; Table 2). Braking and propulsive impulses both increased with faster running velocities (both p < 0.001). Peak braking and propulsive forces paralleled those changes (all p < 0.001). Peak vertical force was independent of horizontal impeding force (p = 0.140), and increased at faster running velocities (p < 0.001). Across running velocities, the changes in braking and propulsive impulses with greater impeding force were correlated (r = -0.97; p < 0.001) indicating that runners who overcame the horizontal impeding forces without reducing their braking impulses substantially, increased their propulsive impulses to a larger extent. However, these respective changes in braking and propulsive impulses were not related to individual changes in metabolic power (p = 0.554 and p = 0.640, respectively).

**Table 1.** Temporal kinematic data for different horizontal impeding force (HIF) conditions.

| Running Velocity (km/h) | HIF (N) | Step Frequency (Hz) | Contact Time (s) | Duty Factor |
| --- | --- | --- | --- | --- |
| 12 | 0 | $2.91 \pm 0.12$ | $0.228 \pm 0.011$ | $0.33 \pm 0.02$ |
| | -4 | $2.91 \pm 0.12$ | $0.228 \pm 0.012$ | $0.33 \pm 0.02$ |
| | -8 | $2.92 \pm 0.11$ | $0.227 \pm 0.012$ | $0.33 \pm 0.02$ |
| 14 | 0 | $2.95 \pm 0.12$ | $0.210 \pm 0.010$ | $0.31 \pm 0.02$ |
| | -4 | $2.95 \pm 0.11$ | $0.210 \pm 0.010$ | $0.31 \pm 0.02$ |
| | -8 | $2.96 \pm 0.09$ | $0.210 \pm 0.010$ | $0.31 \pm 0.02$ |
| 16 | 0 | $2.98 \pm 0.10$ | $0.193 \pm 0.009$ | $0.29 \pm 0.02$ |
| | -4 | $2.99 \pm 0.10$ | $0.192 \pm 0.009$ | $0.29 \pm 0.02$ |
| | -8 | $3.01 \pm 0.09$ | $0.192 \pm 0.010$ | $0.29 \pm 0.02$ |

**Table 2.**
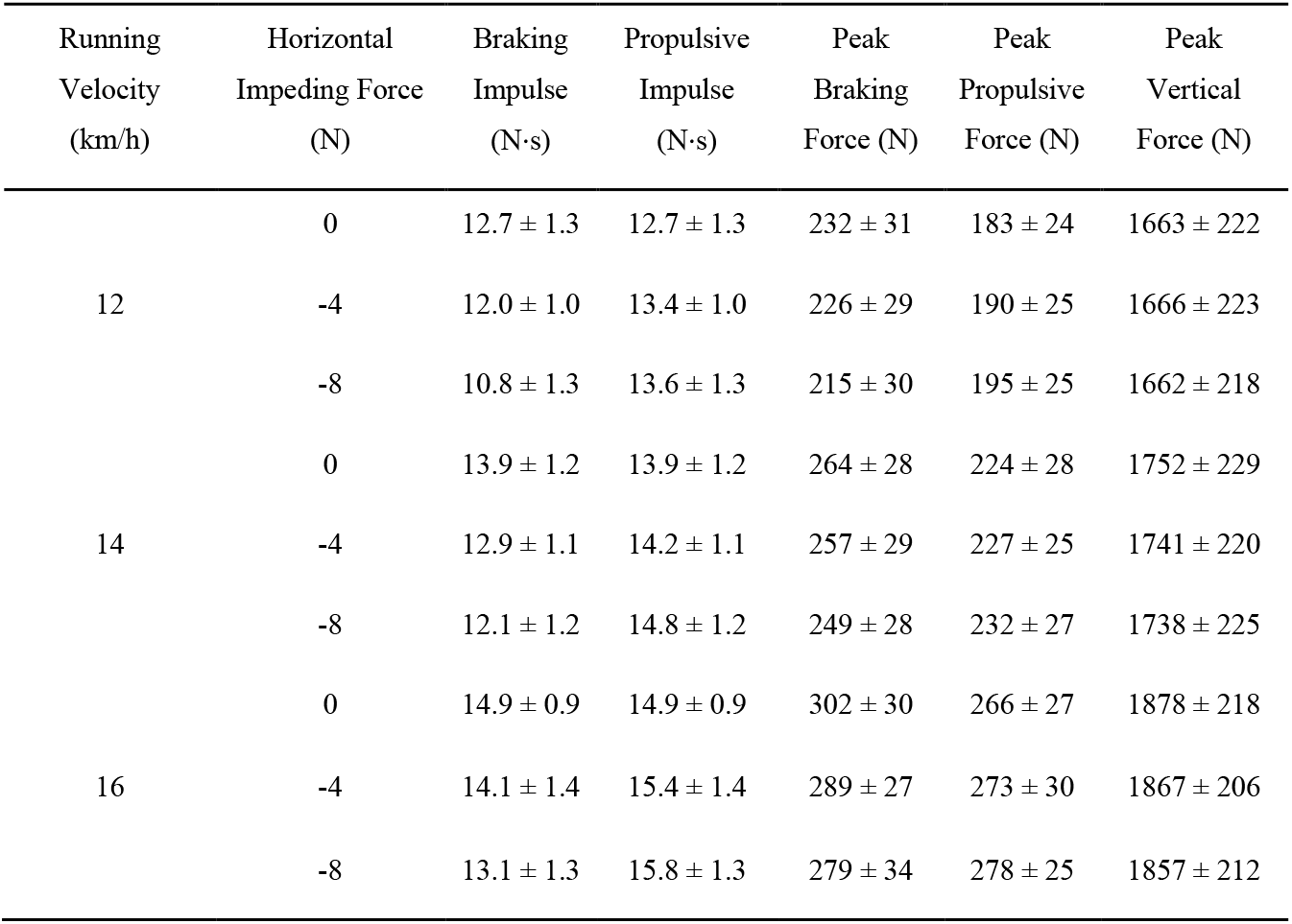
Kinetic data for three running velocities (12, 14 and 16 km/h) and with different horizontal impeding forces (0, 4 and 8 N).

## DISCUSSION

The purpose of our study was to quantify how small resistive forces, similar in magnitude to aerodynamic forces, affect the metabolic power required to run across a range of running velocities. We applied horizontal impeding forces of 0, 4 and 8 N while 12 competitive male runners ran at 12, 14 and 16 km/h. On average, metabolic power increased by 6.13% per 1% BW of horizontal impeding force, but with substantial inter-individual differences, whereby values ranged from 4.75 to 8.14%.

Our results are well within the range of previous studies using horizontal impeding forces during running which span from 4.7 to 7.9% increase per 1% BW (14,15,16). For an air drag force of 1.52% BW at 5.86 m/s, our data indicates a 9.32% increase in metabolic power, slightly higher than the 7% (at 5 m/s) and very close to the 9.2% (at 6.0 m/s) increases in oxygen uptake reported by Jones & Doust (13) and Pugh (2), respectively. Additionally, our data indicates that 8.52% of the metabolic power required during overground running at 4.44 m/s is devoted to overcoming aerodynamic drag, which is close to wind tunnel results from Pugh (8) who calculated 7.5% at 6.0 m/s (extrapolated from observations at 4.47 m/s), but substantially higher than the 4% at 6 m/s that Davies (10) reported based on experiments in the same wind tunnel as Pugh.

In the present study, with greater horizontal impeding force, we found that braking impulses decreased and propulsive impulses increased. Peak braking and propulsive forces paralleled those changes. In addition, there were no effects of horizontal impeding force on step frequency, contact time or duty factor. These results are line with Chang & Kram (16) who evaluated the effect of horizontal impeding forces (0, 3 and 6% of BW) at 3.3 m/s on oxygen uptake and ground reaction forces in well-trained recreational runners. They found the same relation between horizontal impeding forces and braking and propulsive impulses, and between peak braking and propulsive forces, without effects on peak vertical forces, stride frequency, contact time or duty factor.

### Mechanism: force, work or both?

Metabolic power during running is proportional to the vertical force produced to support body weight (20,21,22), with reductions of 1% BW resulting in 0.74% reductions in metabolic power (23) (Teunissen *et al*. 2007). Our data indicates that exerting additional horizontal forces on the ground is ∼8 times costlier than generating vertical forces. This dramatic difference in costs can be explained by the relatively small external moment arms of the vertical ground reaction force component around the leg joints compared to the large external moment arm of the horizontal ground reaction force component around the hip joint (24) (Helseth *et al*. 2008). Larger joint moments require larger muscle forces and thus greater metabolic cost.

Alternatively, from the mechanical work perspective, our apparent efficiency values at first seem paradoxically high as compared to the efficiencies of cycling and concentric muscle contractions. Substantial variations in apparent mechanical efficiency have been reported in the literature for running, mainly related to different methods of mechanical power calculation, muscle efficiency (relation between phosphorylation and contraction coupling) and baseline assumptions for energy cost and elastic energy storage (25,26,27). We did not find systematic effects of horizontal impending force or running velocity on apparent efficiency with values ranging from 40 to 55%. Our results are in line with Bijker *et al*. (28) who found 46% apparent efficiency with extra mechanical power up to 120 W at a running velocity of 8 km/h and to Asmussen & Bonde-Petersen (29) who found 54% apparent efficiency at a running velocity of 10 km/h and extra mechanical power of 69.7 W. However, Lloyd & Zacks (14) found a lower value of apparent efficiency (36%) during 13 km/h running and extra mechanical power up 190 W, which is close to Zacks (15) who reported 39% apparent efficiency for running velocities between 14 to 17 km/h and external mechanical power of 46 to 61 W. Still, all these apparent efficiency values are substantially greater than in cycling (28) and concentric muscle contractions (30,31).

One mechanism behind these enigmatic high apparent efficiency values is the reduction in wasted impulse (32,33). When a person runs on a treadmill at a constant average velocity, they exert equal horizontal braking and propulsive impulses on the ground. The propulsive impulse required to compensate for the braking impulse may be considered “wasted”. In uphill running, braking impulses decrease (34) and thus smaller propulsive impulses are needed to compensate for these smaller braking impulses, i.e. less wasted impulse. Hoogkamer *et al*. (33) reasoned that the change in metabolic power from level to uphill running underestimates the actual metabolic power required to generate the external mechanical power. The actual metabolic power required to generate the external mechanical power is higher, but this is camouflaged by the reduction in the metabolic power related to the reduced wasted impulse with incline that happens simultaneously and is not covered by quantifying the change in metabolic power between level and uphill running. Therefore, apparent efficiency values are higher than the ∼25% efficiency of muscle shortening contractions (30). Future research should address if this concept can also explain the high efficiency values found for running against horizontal impeding forces.

A final issue we wish to address is the 63% efficiency assumed by Polidori *et al*. (7) for converting aerodynamic power to metabolic power. That value comes from Cavagna & Kaneko (35) for the efficiency of performing mechanical work to lift the body center of mass during level running. As Cavagna & Kaneko realized, the discrepancy between 63% and 25% for muscle shortening efficiency reflects that the tendons of the leg muscles act like springs that recycle gravitational potential energy, kinetic energy and elastic energy. In contrast, the work done to overcome aerodynamic forces is dissipated and cannot be elastically recycled.

### Limitations

Our study has some limitations worthy of mention. We applied horizontal impeding forces at the waist (near the center of mass), but air resistive forces during running are spread out over the body. The different shapes of body segments can produce different drag coefficients in specific areas of the body and therefore drag forces vary between segments (36). Future wind tunnel studies are needed to fully validate our findings. In addition, we used fixed horizontal impeding forces (4 and 8 N) across subjects, which represent an average of 0.62 and 1.23% of BW for our subjects (respectively), but ranged from 0.57% to 0.72% BW for 4 N and from 1.15% to 1.44% BW for 8 N. Finally, for our calculations, we assumed that the relative air velocity was equal to the running velocity, but even on calm days, air is never perfectly stationary. Related to this, future studies should evaluate the metabolic cost of running with different wind directions, such as crosswind and tailwind effects, either using an experiment set-up similar to ours or when feasible in a wind tunnel.

## CONCLUSION

We found that metabolic power increase by 6.13% per 1% BW of horizontal impeding force. The increase in metabolic cost results from the cost of generating greater horizontal propulsive forces despite the reduction in wasted impulse.

## Acknowledgements

We thank Clarissa Whiting, Shalaya Kipp, Christian Carmack, Tripp Hurt and Randy Hutchison for their help with data collection.

## Data availability

All data discussed in this manuscript are provided in the tables.

## Authors’ contributions

Conceptualization: Edson Soares da Silva, Rodger Kram, Wouter Hoogkamer; Data collection: Edson Soares da Silva, Wouter Hoogkamer; Data processing: Edson Soares da Silva, Wouter Hoogkamer; Writing - original draft preparation: Edson Soares da Silva, Rodger Kram, Wouter Hoogkamer; Writing - review and editing: Edson Soares da Silva, Rodger Kram, Wouter Hoogkamer.

## Appendix S1

This Appendix addresses several topics. First, we explain in detail how we derived the aerodynamic drag force values from the previous articles which we presented in the Introduction section of this paper. Second, we discuss in more detail how previous articles converted mechanical power estimates to metabolic energy savings.

In a classic study, Hill (1) measured the air resistance forces acting on a scaled physical model of a runner (0.2 m tall) in a small wind tunnel and provided generalized equations for the aerodynamic drag force using only the runner’s height (H) and velocity (v) as inputs. Hill provided the formula 0.15 H^2^ for frontal area (A_f_). Using Kipchoge’s height of 1.67 m yields a frontal area of 0.418 m^2^. The standard equation for aerodynamic drag force (F) (2) in N is:

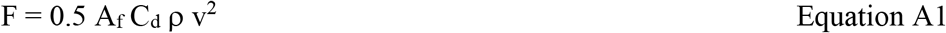

Hill used a C_d_ (coefficient of drag) of 0.9 and air density (ρ) of 1.247 kg/m^3^. Applying Hill’s equation to Kipchoge running solo at a velocity (v) of 5.86 m/s (2-hr marathon pace) yields a force of 8.06 N. Beves & Ferguson (3) used computational fluid dynamic (CFD) modelling to estimate the force acting on Kipchoge running solo at 5.86 m/s as 6.6 N but they did not provide the details behind that value and their simulated depiction of Kipchoge was unrealistically corpulent. Polidori *et al*. (4) used CFD to calculate the air resistance acting on Kenenisa Bekele (the second fastest marathoner to date) running solo at 5.75 m/s. They determined a frontal area of 0.475 m^2^ and used a C_d_ of 0.812 and an air density of 1.219 kg/m^3^. Bekele is slightly shorter and heavier (1.65 m and 56 kg) than Kipchoge. Thus, Polidori *et al*. (4) calculated an aerodynamic drag force of 7.77 N for Bekele.

All three studies described above converted their similar force values first to external mechanical power to then use mechanical efficiency to calculate metabolic power. Multiplying aerodynamic drag force by the running velocity yields mechanical power. Hill’s (1) equations yield 47.2. W of mechanical power for Kipchoge running solo at 5.86 m/s, Beves & Ferguson (3) calculated 38.6 W for Kipchoge solo at 5.86 m/s and Polidori *et al*. (4) found 44.7 W for Bekele solo at the slightly slower velocity (5.75 m/s).

Each of the three studies used different efficiency values. Efficiency is typically calculated as the mechanical power produced divided by the metabolic power required. Hill estimated that a 72.5 kg runner “at longer distances” (unspecified velocity) would have an oxygen uptake of ∼4L O_2_/min (∼55 mlO_2_/kg/min) to provide metabolic power for all of the physiological processes involved in running. That is considerably lower than the ∼70 ml/kg/min of modern, world-class marathoners (5). Hill converted that oxygen uptake of ∼4L O_2_/min to metabolic power assuming exclusively glycogen as the fuel substrate, arriving a value of 1459 W. Hill then divided 46 W of *mechanical* power for just aerodynamic power by the 1459 W of total *metabolic* power required (implicitly, incorrectly assuming an apparent mechanical efficiency of 100%) and concluded that overcoming air resistance comprises only ∼3% of the total metabolic power.

Beves & Ferguson’s (3) model found that when optimally drafting, Kipchoge only needed to produce 10.5 W of mechanical power to overcome drag (a reduction of 28.1 W from solo). Beves & Ferguson then used a value of 300 W for Kipchoge’s sustainable mechanical power which was based on a blogger who used typical values for bicycling. Beves & Ferguson divided the reduction of 28.1 W of mechanical power due to drafting by 300 W of total (cycling) mechanical power yielding a 9 to 10% improvement in running performance compared to running solo. Clearly it is inappropriate to apply a value for cycling to a running.

Polidori *et al*. (4) took yet another approach. They began with estimates for the total mechanical power requirement of 899.6 W when running solo (from an equation from Cavagna & Kaneko (6) combined with their CFD simulation results) and 874.0 W in the optimal drafting configuration. They then used a 63% value for human running efficiency (6) and arrived at 2.8% savings in metabolic power possible with optimal drafting. However, the 63% efficiency value is probably high, in part because it ignores the importance of elastic energy storage and recovery from the tendons in human running (7).

Regardless of the details, the Hill (1), Beves & Ferguson (3) and Polidori *et al*. (4) approaches are intrinsically flawed because the metabolic cost of level running is predominantly determined by muscular force (8,9) and not mechanical power (Heglund *et al*. 1982).

## Appendix S2

### Relative increase in metabolic power per % BW

To be able to compare the findings of Lloyd & Zacks (11), Zacks (12) and Chang & Kram (13), we calculated the relative increases in metabolic power per % BW of resistive force. For Lloyd & Zacks (11) and Zacks (12), we used values for apparent efficiency, metabolic power during “zero-load” running and running velocity reported in Table 2 of each study (as LRE, E_k_ and mean speed, respectively). For Chang & Kram (13), we converted oxygen uptake to metabolic power.

These calculations were straight forward for Chang & Kram (13). They reported data for 8 well-trained recreational runners (5 men and 3 women; 65.8 ± 9.3 kg) that ran at a fixed velocity of 3.3 m/s (11.9 km/h) with horizontal impeding forces of 3 and 6 % BW. We converted oxygen uptake expressed in ml/kg/min into metabolic power (W/kg) by multiplying the average values of oxygen uptake for each condition by 20.9 J/ml oxygen and dividing by 60 seconds/min. Based on the changes in metabolic rate for horizontal impeding forces of 3 and 6 % BW we calculated an average relative increase of 4.7% in metabolic power per 1% BW.

Lloyd & Zacks (11) reported data of 3 male well trained cross-country and track athletes (57.2 ± 0.9 kg) who ran at speeds up to 13 km/h with horizontal impeding forces ranging from 2.2 to 9.6 percent body weight (12.2 to 54.0 N). Zacks (12) reported data for 3 athletes (62.3 ± 9.7 kg (2 athletes participated in both studies)) who ran at velocities ranging from 14 to 17 km/h with horizontal impeding forces ranging from 1.6 to 2.6 percent body weight (9.8 to 15.7 N). Each athlete ran at several different velocities, and with multiple loads at each speed. For each speed, the runner’s metabolic cost during running without resistive forces (E_k_ in kcal/kg/km) and the average apparent efficiency (%) were reported.

First, we converted E_k_ to J/kg/km (factor of 4184 J/kcal). Then, we calculated metabolic power in W/kg by multiplying E_k_ in J/kg/km by the running speed in km/s. Next, we set out to determine the resistive force in % BW and the external mechanical power in W/kg, but Lloyd & Zacks (11) did not provide detailed information about their hanging mass conditions. Instead, we assumed the reported external mechanical power of 70 W for all speeds and calculated the hanging mass that would provide that at each velocity. Zacks (12) states that “At speeds of 14 and 17 km/h the maximum loads were about 1.6 and 1 kg respectively”. We assumed a linear relation between velocity and hanging mass to determine the maximum loads at the other speeds. For each velocity, we then calculated the external mechanical power.

Based on the maximum external mechanical power and the reported average apparent efficiency, we calculated the increase in metabolic power beyond unloaded running. Next, we calculated the relative increase in metabolic power. Finally, we calculated the relative increase in metabolic power per % BW of resistive force. For both studies these calculations resulted in an average relative increase of 7.9% in metabolic power per 1% BW.

## Appendix S3

### Supplementary Table

**Table S1.** Mean metabolic power (W/kg) data for different horizontal impeding forces at the three velocities for each of the 12 subjects tested in the present study.

|  | 12 km/h |  |  | 14 km/h |  |  | 16 km/h |  |  |
| --- | --- | --- | --- | --- | --- | --- | --- | --- | --- |
|  | 0 N | -4 N | -8 N | 0N | -4 N | -8 N | 0 N | -4 N | -8 N |
| 1 | 13.05 | 13.39 | 13.88 | 15.03 | 15.35 | 15.99 | 17.55 | 18.32 | 18.92 |
| 2 | 12.22 | 12.56 | 12.88 | 14.48 | 15.18 | 15.62 | 17.66 | 18.12 | 18.90 |
| 3 | 11.19 | 12.02 | 12.24 | 13.77 | 14.88 | 15.21 | 16.22 | 16.71 | 17.79 |
| 4 | 12.65 | 13.05 | 13.61 | 14.97 | 15.41 | 15.84 | 17.26 | 17.80 | 18.77 |
| 5 | 12.54 | 13.25 | 13.57 | 14.99 | 15.36 | 15.69 | 17.79 | 18.50 | 19.16 |
| 6 | 11.27 | 11.72 | 12.07 | 13.59 | 14.16 | 14.62 | 16.11 | 16.81 | 17.46 |
| 7 | 12.93 | 13.41 | 13.73 | 15.54 | 15.86 | 16.44 | 18.19 | 18.46 | 19.08 |
| 8 | - | - | - | - | - | - | 16.25 | 16.90 | 17.90 |
| 9 | 11.64 | 12.19 | 12.51 | 14.36 | 15.09 | 15.77 | 16.69 | 17.39 | 18.51 |
| 10 | 12.02 | 12.38 | 12.89 | 15.02 | 15.28 | 16.05 | 17.17 | 17.75 | 18.34 |
| 11 | 11.30 | 11.76 | 12.07 | 15.39 | 15.68 | 16.30 | 16.45 | 16.87 | 17.65 |
| 12 | 10.86 | 11.36 | 11.92 | 13.37 | 13.88 | 14.57 | 16.95 | 17.56 | 18.36 |

